# Divergence of alternative sugar preferences through modulation of the expression and activity of the Gal3 sensor in yeast

**DOI:** 10.1101/2023.01.12.523712

**Authors:** Josep Fita-Torró, Krishna B. S. Swamy, Amparo Pascual-Ahuir, Markus Proft

**Affiliations:** Department of Molecular and Cellular Pathology and Therapy, Instituto de Biomedicina de Valencia IBV-CSIC, 46010 Valencia, Spain; Division of Biological and Life Sciences, School of Arts and Sciences, Ahmedabad University, Ahmedabad, 380009, India; Department of Biotechnology, Instituto de Biología Molecular y Celular de Plantas, Universitat Politècnica de València, 46022 Valencia, Spain

**Author notes:** Correspondence (AP-A); (MP).

**Keywords:** Budding yeast, sugar utilization, *GAL* switch, diauxic growth, *GAL* evolution, *GAL1*, *GAL3*, *Saccharomyces bayanus*

## Abstract

Optimized nutrient utilization is crucial for the progression of microorganisms in competing communities. Here we investigate how different budding yeast species and ecological isolates have established divergent preferences for two alternative sugar substrates: Glucose, which is fermented preferentially by yeast, and galactose, which is alternatively used upon induction of the relevant *GAL* metabolic genes. We quantified the dose-dependent induction of the *GAL1* gene encoding the central galactokinase enzyme, and found that a very large diversification exists between different yeast ecotypes and species. The sensitivity of *GAL1* induction correlates with the growth performance of the respective yeasts with the alternative sugar. We further define some of the mechanisms, which have established different glucose/galactose consumption strategies in representative yeast strains by modulating the activity of the Gal3 inducer. (1) Optimal galactose consumers, such as *Saccharomyces bayanus*, contain a hyperactive *GAL3* promoter, sustaining highly sensitive *GAL1* expression, which is not further improved upon repetitive galactose encounters. (2) Desensitized galactose consumers, such as *S. cerevisiae* Y12, contain a less sensitive Gal3 sensor, causing a shift of the galactose response towards higher sugar concentrations even in galactose experienced cells. (3) Galactose insensitive sugar consumers, such as *S. cerevisiae* DBVPG6044, contain an interrupted *GAL3* gene, causing extremely reluctant galactose consumption, which however still is improved upon repeated galactose availability. In summary, different yeast strains and natural isolates have evolved galactose utilization strategies, which cover the whole range of possible sensitivities by modulating the expression and/or activity of the inducible galactose sensor Gal3.

## Introduction

Microorganisms have evolved sophisticated metabolic regulation to adapt to environments with continuously changing availability of nutrients. An important example is the usage of different sugar sources by unicellular fungi such as budding yeast (1). *Saccharomyces cerevisiae* and evolutionarily related species have specialized towards the preferential fermentation of glucose over alternative sugar moieties (2). This metabolic behavior is driven by a regulatory network called glucose repression, which inhibits the expression of genes necessary for the uptake and breakdown of alternative carbohydrates in the presence of glucose (2). One alternative sugar, which is frequently found in natural environments is the monosaccharide galactose (3). Budding yeasts can assimilate galactose by the induced expression of three clustered structural genes, *GAL1, GAL7* and *GAL10*, whose enzymatic products are responsible for converting galactose into glucose-1-phosphate in order to generate energy for the cell via glycolysis (4). The galactose-utilization (*GAL*) pathway in *Saccharomyces cerevisiae* has been established as a model system to understand the function and evolution of eukaryotic metabolic regulation and to unravel the principles of microbial decision-making (5–7).

*GAL* gene expression in *S. cerevisiae* is strictly repressed in the presence of glucose (8, 9), and massively induced when glucose levels decline and galactose is available (10). The Zn-cluster transcription factor Gal4 is the master activator of *GAL* genes binding to a highly enriched recognition site with the consensus sequence 5’-CGG-N_11_-CCG-3’ (11–13). However, in the absence of galactose, Gal4 is kept in an inactive state by direct binding of the Gal80 repressor protein (14, 15). This inhibition is counteracted by the additional binding of the Gal3 sensor upon its activation by galactose (16, 17). Thus, Gal3 operates the *GAL* on/off switch in a galactose concentration-dependent manner (18). Since the expression of *GAL3* itself is inducible by galactose and Gal4, the *GAL* switch acquires additional modes of plasticity, which might fine-tune the sensitivity and efficiency of the *GAL* response during the adaptation to different galactose availability (19, 20). Indeed, at certain galactose limiting conditions and dependent on the carbon source history and the genetic background, the transition from the uninduced (off) to the induced (on) state is not uniform within a yeast population, which leads to the co-existence of responders and non-responders or bimodal response (19, 21, 22). It has been very recently shown that the levels of the Gal3 sensor, either artificially modulated or dictated by different natural alleles, dominantly mediates the modality of the *GAL* switch (23). The constitutive overexpression of Gal3 leads to a galactose independent activation of the *GAL* response, highlighting the dominant function of the sensor in the control of *GAL* gene induction (24).

Another important feature of the *S. cerevisiae GAL* response, which accounts for its environmental adaptability, is its enhanced activation after previous galactose encounter or transcriptional memory (25, 26). In naïve yeast cells, *GAL* gene induction is slow and inefficient especially at low galactose concentrations. After a previous galactose encounter, transcriptional activation of *GAL* is faster and sensitive to low inducer concentrations, and this memory state is maintained during several generations (27–29). Enhanced *GAL* induction during memory depends on the accumulation of the Gal1 galactokinase or the Gal3 galactose sensor (28, 30, 31). Both proteins recognize galactose and associate with the Gal80 repressor, thus switching Gal4 to the active form and unleashing *GAL* gene expression (18, 32). Faster re-activation of the *GAL* response might be advantageous in environments with frequent galactose availability. Induction of Gal1/Gal3 in the previous galactose encounter is a way to promote faster adaptation to the alternative sugar, and accordingly, the Gal1 or Gal3 expression levels have been found to correlate with the efficiency of *GAL* memory.

In this work, we investigate to what degree different galactose recognition and consumption behaviors have been evolved in natural *S. cerevisiae* isolates from different ecological niches and evolutionarily related species such as *S. bayanus, S. mikatae* and *S. paradoxus*. We found that across 12 different species a very broad spectrum of galactose preferences exists, from almost complete galactose reluctance to highly sensitive *GAL* responses. These behaviors correlate with divergent dose-response profiles of *GAL1* induction and have consequences for glucose/galactose growth performance, diauxic shift, galactose memory and susceptibility to toxic sugar analogues. We further characterize representative galactose consumption strategies, which have evolved by tuning Gal3 expression/activity: the ultra-sensitive galactose response of *S. bayanus*, the specialization towards higher galactose concentrations of *S. cerevisiae* sake strain Y12, and galactose insensitivity in West African *S. cerevisiae* strains.

## Materials and methods

### Yeast strains and growth conditions

The yeast strains used in this work are described in Table 1. Yeast cultures were grown at 28°C in Yeast Extract Peptone Dextrose (YPD) or Galactose (YPGal) media containing 2% glucose or galactose or indicated mixtures of both sugars, or in Synthetic Dextrose (SD), Raffinose (SRaf) or Galactose (SGal) media containing 0.67% yeast nitrogen base with ammonium sulfate and without amino acids, 50mM succinic acid (pH 5.5) and 2% of the respective sugar. According to the auxotrophies of each strain, methionine (10 mg/l), histidine (10 mg/l), leucine (10 mg/l) or uracil (25 mg/l) were supplemented. Yeast cells were transformed by the lithium acetate/PEG method described by (33). The glucose analogue glucosamine hydrochloride (GlcN, Sigma Aldrich) was added to the growth medium at the indicated final concentrations from a 10% stock solution in H_2_O.

**Table 1:** Yeast strains used in this study

| Strain | Genotype | Source |
| --- | --- | --- |
| **S. cerevisiae BY4741** | MATa; *his3*Δ1; *leu2*Δ0; *met15*Δ0; *ura3*Δ0 | EUROSCARF |
| **S. paradoxus** | ho::HPH; *ura3*; *lys2*; *his3*Δ::3xHA; *leu2*Δ::3xHA | Jun-Yi Leu |
| **S. mikatae** | ho::HPH; *ura3*; *lys2*; *his3*Δ::3xHA; *leu2*Δ::3xHA | Jun-Yi Leu |
| **S. bayanus** | ho::HPH; *ura3*; *his3*Δ::3xHA; *leu2*Δ::3xHA | Jun-Yi Leu |
| **S. cerevisiae UWOPS05-217.3** | ho::HGB; *ura3*::G418-barcode; *his3*::clonat | Jun-Yi Leu |
| **S. cerevisiae NCYC110** | ho::HGB; *ura3*::G418-barcode; *his3*::clonat | Jun-Yi Leu |
| **S. cerevisiae DBVPG6044** | ho::HGB; *ura3*::G418-barcode; *his3*::clonat | Jun-Yi Leu |
| **S. cerevisiae Y12** | ho::HGB; *ura3*::G418-barcode; *his3*::clonat | Jun-Yi Leu |
| **S. cerevisiae YJM981** | ho::HGB; *ura3*::G418-barcode; *his3*::clonat | Jun-Yi Leu |
| **S. cerevisiae L1374** | ho::HGB; *ura3*::G418-barcode; *his3*::clonat | Jun-Yi Leu |
| **S. cerevisiae YPS128** | ho::HGB; *ura3*::G418-barcode; *his3*::clonat | Jun-Yi Leu |
| **S. cerevisiae YPS606** | ho::HGB; *ura3*::G418-barcode; *his3*::clonat | Jun-Yi Leu |
| **BY4741 GAL1-lucCP<sup>+</sup>** | BY4741 with plasmid pAG413-GAL1-lucCP<sup>+</sup> (HIS3) | This study |
| **S. paradoxus GAL1-lucCP<sup>+</sup>** | *S. paradoxus* with plasmid pAG413-GAL1-lucCP<sup>+</sup> (HIS3) | This study |
| **S. mikatae GAL1-lucCP<sup>+</sup>** | *S. mikatae* with plasmid pAG413-GAL1-lucCP<sup>+</sup> (HIS3) | This study |
| **S. bayanus GAL1-lucCP<sup>+</sup>** | *S. bayanus* with plasmid pAG413-GAL1-lucCP<sup>+</sup> (HIS3) | This study |
| **UWOPS05-217.3 GAL1-lucCP<sup>+</sup>** | UWOPS05-217.3 with plasmid pAG413-GAL1-lucCP<sup>+</sup> (HIS3) | This study |
| **NCYC110 GAL1-lucCP<sup>+</sup>** | NCYC110 with plasmid pAG413-GAL1-lucCP<sup>+</sup> (HIS3) | This study |
| **DBVPG6044 GAL1-lucCP<sup>+</sup>** | DBVPG6044 with plasmid pAG413-GAL1-lucCP<sup>+</sup> (HIS3) | This study |
| **Y12 GAL1-lucCP<sup>+</sup>** | Y12 with plasmid pAG413-GAL1-lucCP<sup>+</sup> (HIS3) | This study |
| **YJM981 GAL1-lucCP<sup>+</sup>** | YJM981 with plasmid pAG413-GAL1-lucCP<sup>+</sup> (HIS3) | This study |
| **L1374 GAL1-lucCP<sup>+</sup>** | L1374 with plasmid pAG413-GAL1-lucCP<sup>+</sup> (HIS3) | This study |
| **YPS128 GAL1-lucCP<sup>+</sup>** | YPS128 with plasmid pAG413-GAL1-lucCP<sup>+</sup> (HIS3) | This study |
| **YPS606 GAL1-lucCP<sup>+</sup>** | YPS606 with plasmid pAG413-GAL1-lucCP<sup>+</sup> (HIS3) | This study |
| **BY4741 GAL3<sub>BY4741</sub>-lucCP<sup>+</sup>** | BY4741 with plasmid pAG423-GAL3<sub>BY4741</sub>-lucCP<sup>+</sup> (HIS3) | This study |
| **BY4741 GAL3<sub>S.bayanus</sub>-lucCP<sup>+</sup>** | BY4741 with plasmid pAG423-GAL3<sub>S.bayanus</sub>-lucCP<sup>+</sup> (HIS3) | This study |
| **BY4741 GAL3<sub>DBVPG6044</sub>-lucCP<sup>+</sup>** | BY4741 with plasmid pAG423-GAL3<sub>DBVPG6044</sub>-lucCP<sup>+</sup> (HIS3) | This study |
| **BY4741 GAL3<sub>Y12</sub>-lucCP<sup>+</sup>** | BY4741 with plasmid pAG423-GAL3<sub>Y12</sub>-lucCP<sup>+</sup> (HIS3) | This study |
| **BY4741 GAL1-GFP** | BY4741 with plasmid pRS416-GAL1<sub>BY4741</sub>-eGFP (URA3) | This study |
| **Y12 GAL1-GFP** | Y12 with plasmid pRS416-GAL1<sub>BY4741</sub>-eGFP (URA3) | This study |
| **BY4741 *gal3*Δ GAL1-lucCP<sup>+</sup>** | BY4741 *gal3::KAN* with plasmid pAG413 GAL1<sub>BY4741</sub>-lucCP<sup>+</sup> (HIS3) | This study |
| **Gal3<sub>BY4741</sub> GAL1-lucCP<sup>+</sup>** | BY4741 *gal3::KAN* GAL1<sub>BY4741</sub>-lucCP<sup>+</sup> with plasmid pAG416-GAL1p-GAL3<sub>BY4741</sub> (URA3) | This study |
| **Gal3<sub>Y12</sub> GAL1-lucCP<sup>+</sup>** | BY4741 *gal3::KAN* GAL1<sub>BY4741</sub>-lucCP<sup>+</sup> with plasmid pAG416-GAL1p-GAL3<sub>Y12</sub> (URA3) | This study |
| **DBVPG6044 GPD** | DBVPG6044 with plasmid pAG416-GPD-*ccdB* (URA3) | This study |
| **NCYC110 GPD** | NCYC110 with plasmid pAG416-GPD-*ccdB* (URA3) | This study |
| **DBVPG6044 Gal3<sub>BY4741</sub>** | DBVPG6044 with plasmid pAG416-GPD-GAL3<sub>BY4741</sub> (URA3) | This study |
| **NCYC110 Gal3<sub>BY4741</sub>** | NCYC110 with plasmid pAG416-GPD-GAL3<sub>BY4741</sub> (URA3) | This study |

### Plasmid constructions

The single-copy destabilized luciferase reporter plasmid pAG413-GAL1_BY4741_-lucCP_+_ (34) was used to quantify the *GAL1* dose response of different yeast isolates. Multi-copy reporter plasmids expressing destabilized luciferase under control of different *GAL3* promoter alleles were constructed by insertion of PCR amplified fragments containing the 900bp upstream of the ATG into luciferase vector pAG423-lucCP_+_ (34). For the quantification of *GAL1* activation by flow cytometry, a single-copy GFP reporter was constructed by insertion of the *GAL1*_BY4741_ upstream regulatory region in plasmid pRS416-GFP (35). For the quantitative analysis of the Gal3 sensors of BY4741 and Y12, we cloned the respective *GAL3* ORF regions in the Gateway single copy expression vector pAG416-GAL1-ccdB (36). Complementation of yeast cells with the wild type BY4741 copy of *GAL3* was done using the single copy expression plasmid pAG416-GPD-GAL3_BY4741_ (28). Primers used for plasmid constructions are summarized in Table 2. All *GAL3* promoter and ORF variants used in this study were sequenced for verification purposes and for the identification of specific nucleotide changes occurring in the natural yeast variants.

**Table 2:**
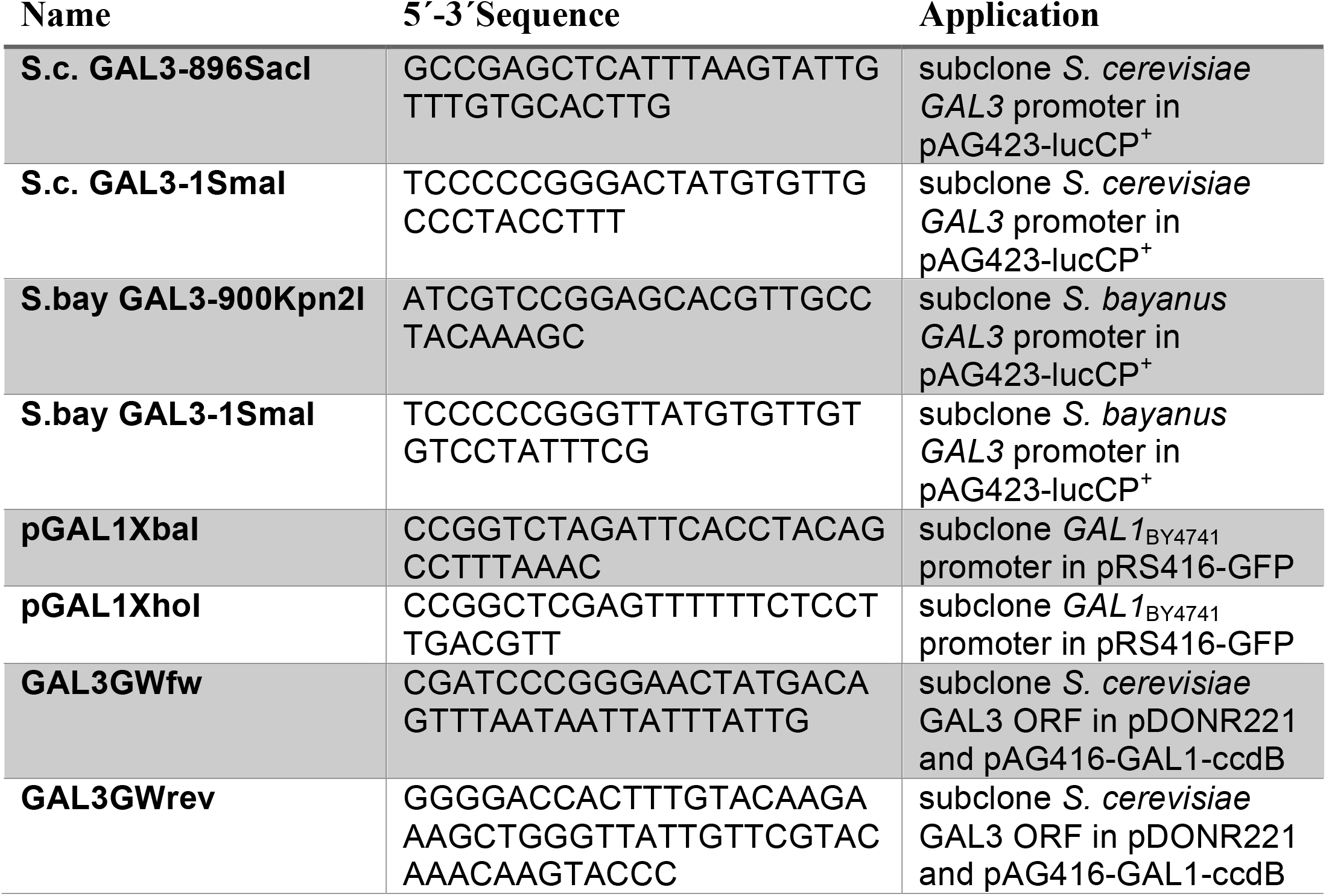
Oligonucleotide primers used in this study

### Live-cell luciferase assays

Yeast strains containing the indicated luciferase fusion genes were grown at 28°C in synthetic raffinose (SRaff) medium lacking histidine at pH=3 to low exponential phase (OD600 = 1-2). Culture aliquots were then incubated with 0.5mM luciferin (free acid, Synchem, Germany) on a roller at 28°C for 1h. The cells were then transferred in 135μl aliquots to white 96 well plates (Nunc) containing the indicated concentrations of galactose. The light emission was continuously recorded on a GloMax microplate luminometer (Promega) in at least 3 biological replicas. Data were processed with Microsoft Excel software. For representation of the relative light units, we normalized the raw data for the number of cells in each individual assay. The maximal synthesis rate was calculated as described previously (34).

### Quantitative growth assays

For the quantitative estimation of growth parameters, fresh overnight precultures of the indicated yeast isolates in YPD or YPGal media were diluted in triplicate in the assay medium in multiwell plates to a starting OD600 = 0.1. Growth was then constantly monitored at 28°C on a Tecan Spark multiplate reader for the indicated times. The growth curves were processed in Microsoft Excel and absolute growth ratios and lag phases calculated.

### Transcriptional memory experiments

For memory experiments at *GAL1*, cells containing the GAL1p-lucCP^+^ reporter gene were grown over night in synthetic Raffinose (2%) medium lacking histidine to exponential growth phase. A first round of induction was then performed for 2 h with 1% galactose, while naïve cells remained in SRaf medium. Both cell cutures were then pelleted, washed once with H2O, and then incubated in SD (2% glucose) medium for 1h. Finally, cells were washed again and resuspended in SRaf medium containing 0.5 mM luciferin for 90 min before starting the next induction with the indicated galactose concentrations. Light emission was then monitored continuously as described above for the standard luciferase assays.

### Flow cytometry

Cells harboring a single-copy GAL1p-eGFP reporter were pre-grown in synthetic raffinose medium (SRaf) without uracil to exponential growth phase. Cells were diluted to OD_600_ = 0.1 and induced with the indicated galactose concentrations. Before induction and at the indicated times of galactose supplementation, cell aliquots were passed through a MACS Quant 10 flow cytometer (Miltenyi Biotec). GFP was excited with a 488nm laser and emission was detected applying a 525/550nm band pass filter set. All GAL1-GFP induction experiments were performed on 3 independent biological samples. 20000 cells were analyzed for each time point, and aggregated cells (<5%) were discarded from further analysis. For the detection of the induced single cell fraction, the uninduced fluorescence intensity was defined for the cell population at time point 0, and during the induction process, all cells with higher fluorescence intensities were counted as “positive”. The mean fluorescence intensity was additionally determined in the single cell fraction at each time point of galactose induction.

## Results

### Dose-dependent *GAL1* induction profiles reveal a broad spectrum of different sugar preferences across natural budding yeast strains

Laboratory *Saccharomyces cerevisiae* strains typically show a strictly galactose concentration-dependent activation of *GAL* genes upon the switch from glucose to galactose (28). High galactose inducer concentrations are needed to reach optimal induction kinetics. We therefore reasoned that divergent galactose preferences could be quantitatively determined by comparing the dose dependent induction profiles of *GAL1* across a variety of yeast strains from different ecological origins. In order to obtain truly quantitative gene expression profiles in real time, we employed the extremely unstable lucCP^+^ luciferase derivative in the laboratory reference strain BY4741 (34), as well as in 8 different *S. cerevisiae* natural isolates and in the evolutionarily close *S. paradoxus, S. mikatae* and *S. bayanus* strains (Figure 1A). We recorded the dose-responsive activation profiles for *GAL1* applying an exhaustive range of galactose inducer concentrations (Figure 1B). All strains tested showed a measurable activation of luciferase activity over the initial 2.5 hours of induction, except the West African isolates DBVPG6044 and NCYC110. We noticed important differences when we compared the absolute induction levels along the increasing galactose concentrations for each yeast strain (Figure 1C). *S. mikatae* and especially *S. bayanus* respond to low galactose concentrations much more efficiently and reach optimal induction velocities at much lower galactose concentrations as compared to the BY4741 laboratory strain. Both yeast species also display a prolonged activation of the *GAL1* gene, which in the case of S. bayanus leads to almost 7 fold enhanced induction ratios. This suggests that S. *mikatae* and *S. bayanus* have evolved towards a more sensitive galactose recognition. The natural Y12 *S. cerevisiae* isolate has yet a distinctive pattern of galactose induction. In this case, *GAL1* induction is very efficient at high inducer concentrations, while it is slow and inefficient upon low galactose stimulation. This suggests that *S. cerevisiae* Y12 has adapted to discriminate more different galactose concentrations and recognizes the alternative sugar preferentially at higher doses. Most other natural *S. cerevisiae* isolates tested here showed a *GAL1* dose response, which resembled the BY4741 reference. However, *S. paradoxus* responds to increasing galactose concentrations much more inefficiently and in the DBVPG6044 and NCYC110 isolates, a rapid galactose response was completely absent. These data demonstrate that natural yeast isolates have evolved very different galactose response strategies, whose consequences for growth in the presence of the alternative sugar will be investigated next.

**Fig. 1.**
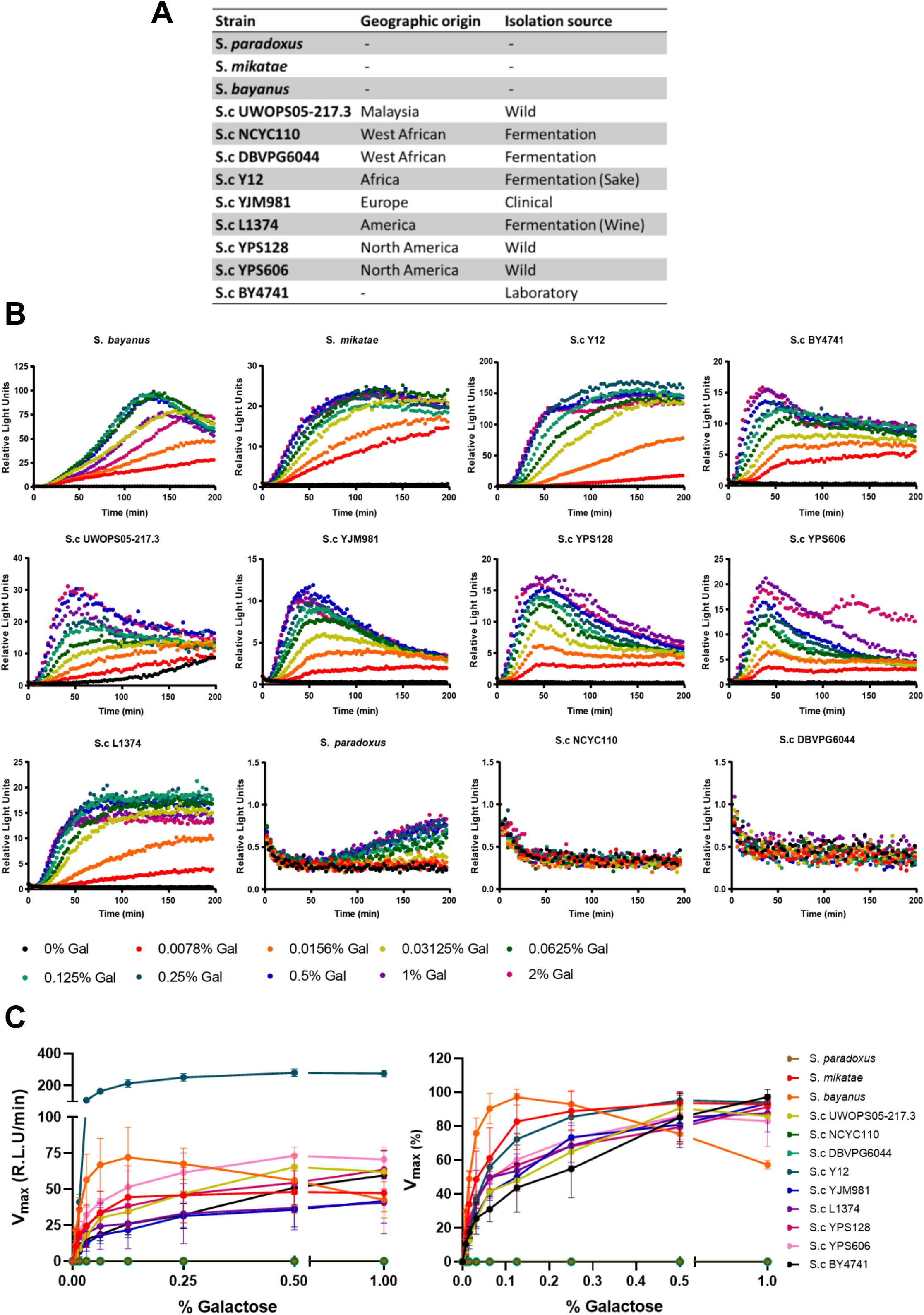
Dose-dependent *GAL1* expression across natural yeast isolates. (A) Yeast strains and natural isolates from different geographical origin used in this study. (B) Time-elapsed study of the dose dependent induction of *GAL1* in the indicated yeast strains. A single copy GAL1-lucCP^+^ reporter was used in continuous live cell luciferase assays. Results are depicted from at least 3 biological replicates. Light emission at time point 0 was set to 1 for all induction curves. (C) Comparison of the sensitivity of *GAL1* induction. Representation of the maximal light increment (v_max_) versus the galactose inducer concentration for each yeast strain. Left panel: Absolute v_max_ values plotted against gal concentrations, right panel: v_max_ values set to 100% for the inducer concentration, which leads to maximal induction.

### Sensitivity of *GAL* induction correlates with growth performance and glucose/galactose selectivity

We compared quantitative growth parameters for all the natural yeast strains included in this study with respect to glucose and galactose utilization. As shown in Figure 2A, the difference of the absolute growth rates for both sugars is extremely divergent among the yeast strains. In the BY4741 reference strain, the growth rate on galactose slows down to 60% as compared to glucose. *S. mikatae* and especially *S. bayanus* demonstrate less sugar discrimination. Remarkably, *S. bayanus* cells grow with indistinguishable rates on both sugar substrates. This correlates well with their better and more sensitive *GAL1* gene induction. On the lower end, we identify yeast strains with a much more pronounced sugar discrimination, which grow on galactose more inefficiently. These strains are the *S. cerevisiae* Y12 isolate, *S. paradoxus* and the DBVPG6044 and NCYC110 isolates, which altogether are characterized by their sub-optimal *GAL1* induction profiles. These growth differences are also well recapitulated when we calculated the lag phase needed by all strains to engage in active growth when switched from glucose to galactose media (Figure 2A). We next tested the performance of selected yeast strains during the diauxic shift on mixed sugar media with limited amounts of glucose (Figure 2B). Under these conditions, the cells have to switch from glucose to galactose utilization. Moderate *GAL1* inducers such as BY4741 or Y12, show a characteristic diauxic lag phase separating glucose and galactose consumption. However, optimized *GAL1* inducers, *S. mikatae* and S. *bayanus*, do not display a diauxic lag and grow continuously on the mixed sugar media. Interestingly, *S. bayanus* shows a slight growth disadvantage on glucose media as compared to BY4741, which is more than compensated during the glucose-galactose switch where S. *bayanus* cells are able to outgrow BY4741 (Figure 2B). As expected, poor *GAL1* inducers DBVPG6044 and NCYC110 show an extreme diauxic lag and low growth rates on galactose media. These data indicated that yeast strains with an optimized *GAL* gene induction acquire a growth advantage during mixed sugar or pure galactose growth.

**Fig. 2.**
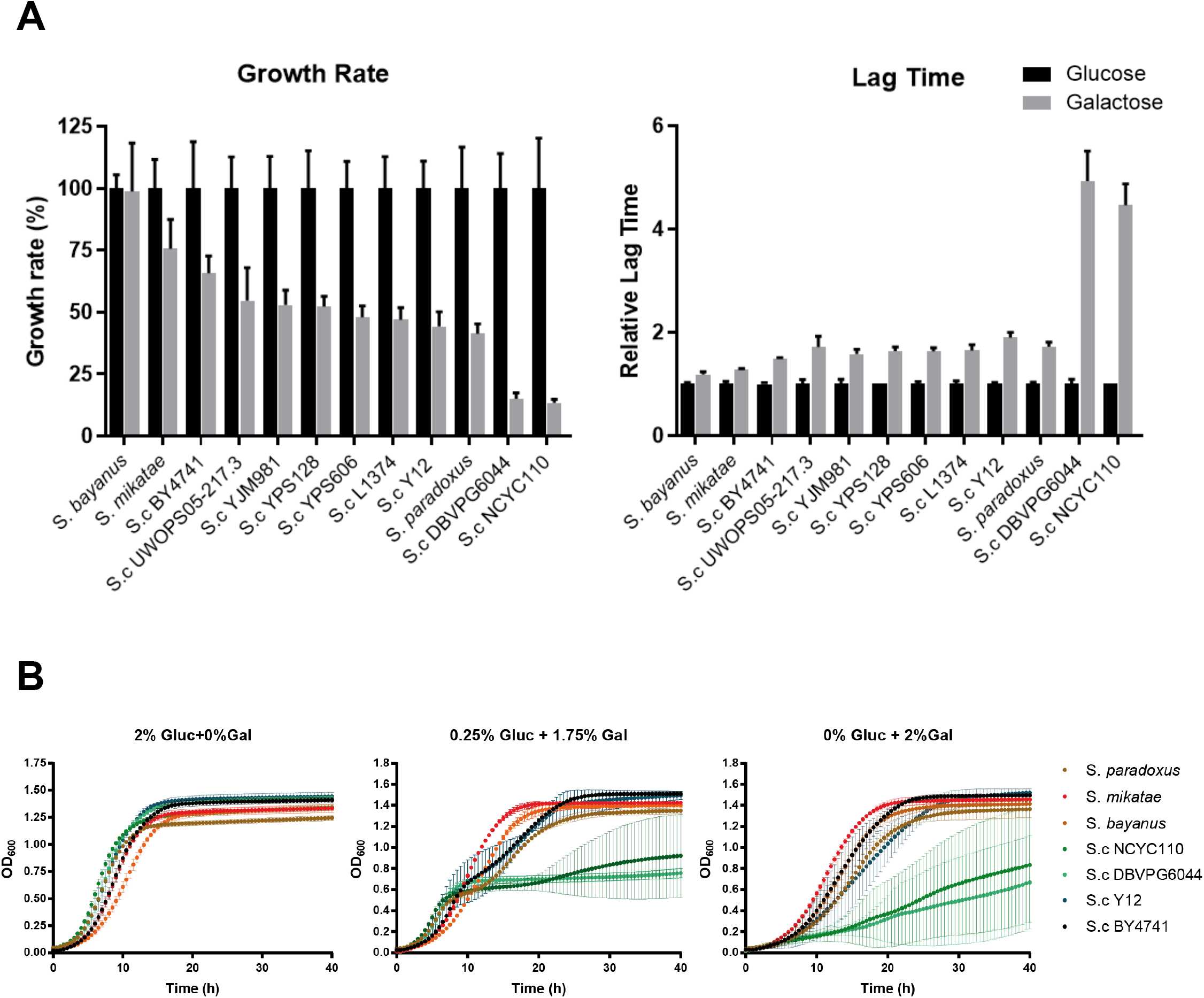
Glucose/galactose growth performance across natural yeast isolates. (A) Left panel: The growth rates on 2% glucose or galactose media were determined for the indicated yeast strains pregrown on glucose medium. The velocity on glucose medium was set to 100% for each strain. Right panel: Representation of the lag time needed for each strain to reach 50% growth on the indicated media. The lag time for glucose growth was set to 1 for each strain. Represented are the mean values (n = 3) +/- SD. (B) Continuous growth curves for the indicated yeast strains on pure glucose or galactose (2%) media, or on a mixture of glucose/galactose (0.25%/1.75%). Data are mean values (n = 3) +/- SD.

We next wanted to know whether the inefficient galactose growth performance of some selected yeast strains could be improved by previous galactose adaptation. We therefore compared the growth parameters for galactose of cells that were either pre-grown in glucose or galactose (Figure 3). For the best galactose performers including *S. mikatae* and S. *bayanus*, but also the BY4741 laboratory strain, galactose pre-growth did not improve neither the growth rate nor the lag phase on galactose media (Figure 3A and 3B). However, poor galactose performers such as *S. cerevisiae* Y12, DBVPG6044 and NCYC110, largely improve their growth rates and cut down the lag phase on galactose media after previous galactose adaptation. This suggests that all strains tested have conserved the ability to metabolize galactose as an alternative energy source, however, signaling differs substantially in different yeasts creating very divergent galactose preferences.

**Fig. 3.**
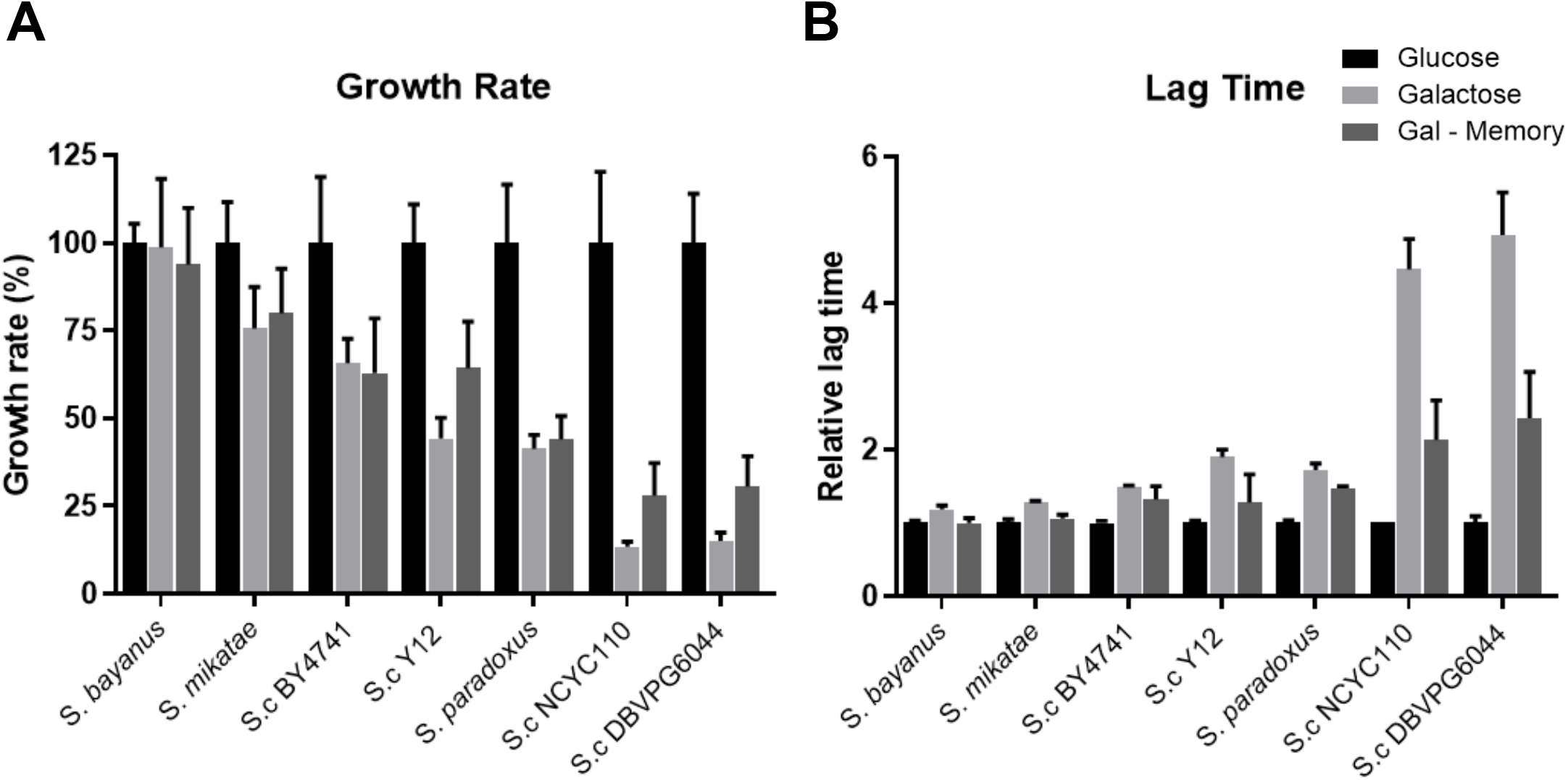
Growth performance after adaptation to galactose medium. The indicated yeast strains were pre-grown in glucose (Glucose, Galactose) or galactose (Gal-Memory) media and then diluted in the indicated growth media. (A) Absolute growth rates and (B) the lag times were compared by setting the glucose growth rate and lag time for each strain to 100% or 1, respectively. Data are mean values (n = 3) +/- SD.

### Specialization towards highly sensitive galactose consumption: the *S. bayanus* case

In our previous experiments, *S. bayanus* was characterized as the most efficient galactose consumer based on its highly sensitive and efficient *GAL1* induction profile, equal growth performance on glucose or galactose media and an optimized transition from glucose to the alternative sugar in mixed sugar environments. A closer inspection of the *GAL1* dose response data revealed that *S. bayanus* displays a remarkably robust gene induction, which is almost complete at very low galactose concentrations (0.03%) (Figure 4A). One consequence of this galactose hypersensitivity is a growth advantage over other yeasts with a more repressed galactose signaling, such as BY4741, with a pronounced adaptation phase during the switch from one sugar to the other (Figure 4B). We reasoned that *S. bayanus* was less dependent on glucose fermentation and more readily metabolized galactose on mixed substrates. We tested this hypothesis by the application of the naturally occurring glucose analogue glucosamine (2-amino-2-deoxy-glucose). Glucosamine is taken up and phosphorylated by *Saccharomyces* species, however, cannot be further metabolized via glycolysis, which results in a strong growth inhibition (37). When BY4741 reference and *S. bayanus* cells were offered mixtures of glucosamine and galactose, we found that *S. bayanus* was much less susceptible to glucosamine growth arrest (Figure 4C). These results confirmed that the *S. bayanus* strain had a superior affinity for galactose consumption, which caused the observed hyperresistance to the glucose analogue. We next addressed the question, whether a previous galactose encounter improved the transcriptional response in these two divergent yeasts (Figure 4D). *S. cerevisiae* laboratory strains such as BY4741 are characterized by a strong improvement of their induction kinetics, both efficiency and sensitivity, by mechanisms of transcriptional memory (28, 31). As expected, a robust transcriptional memory effect was determined for the *S. c*. BY4741 strain. However, *S. bayanus* displayed an already optimal *GAL1* dose response in naïve cells, which was not at all improved after galactose pre-treatment (Figure 4E). These data suggest that *S. bayanus* has evolved an optimized galactose signaling, which is constitutively active and not susceptible to improvement by previous galactose consumption. We next wanted to identify the molecular bases for this galactose specialization.

**Fig. 4.**
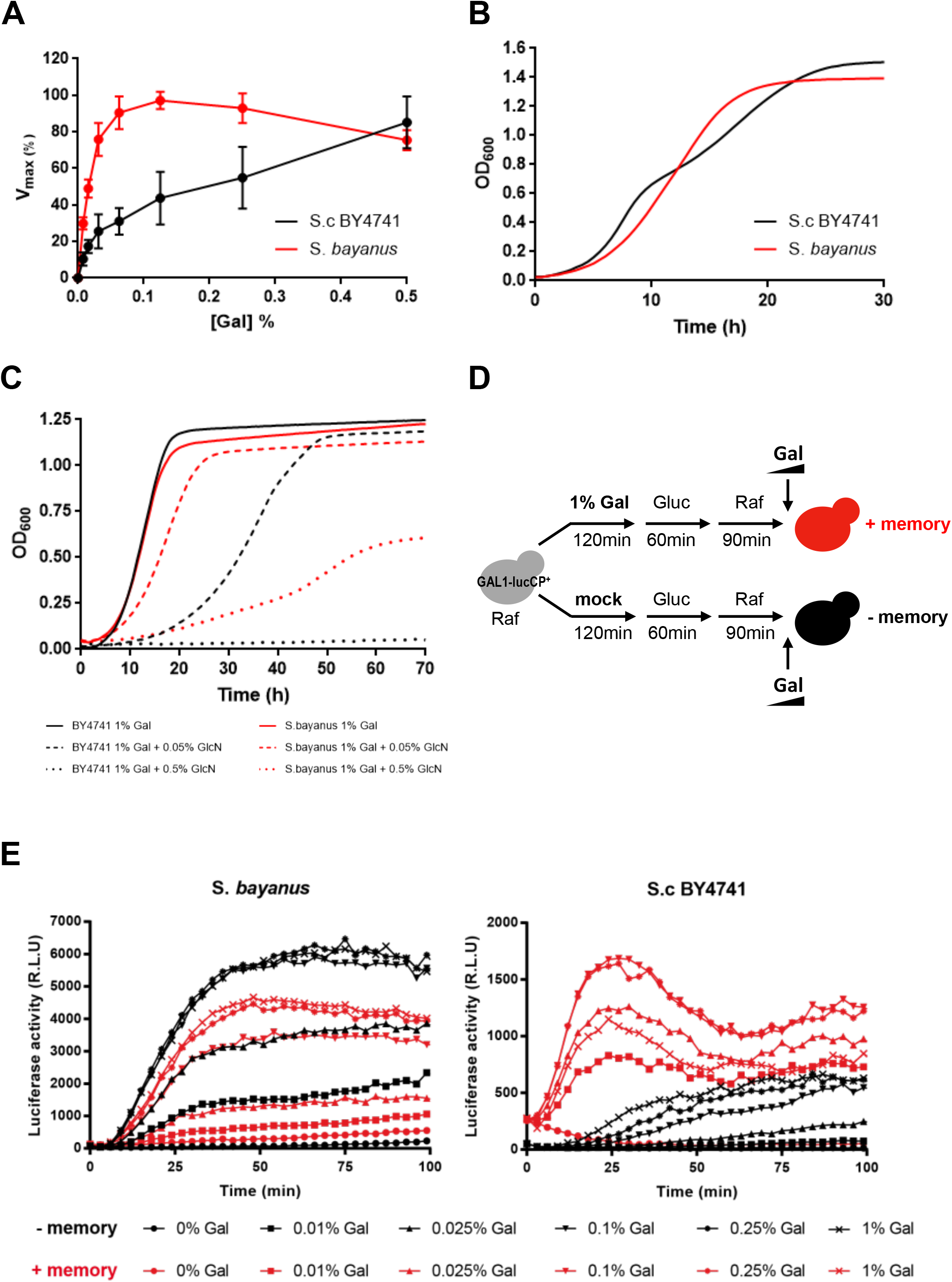
Features of highly sensitive galactose consumption in *Saccharomyces bayanus*. (A) Optimized gene expression induction by very low galactose concentrations. The relative light increment (v_max_) of a *GAL1*-luciferase reporter is compared between *S. cerevisiae* BY4741 and *S. bayanus* for the indicated inducer concentrations. Data are taken from the experiment described in Figure 1B, C. (B) Lack of diauxic growth delay. Yeast strains were grown in a glucose/galactose mixture (0.25%/1.75%). (C) Resistance to the glucose analogue glucosamine. Yeast cells were grown in pure galactose medium (1%) with or without the addition of the indicated amounts of glucosamine (GlcN). (D) Experimental setup for the determination of transcriptional memory upon galactose pre-treatment. (E) *S. bayanus* lacks a transcriptional galactose memory. Comparison of the *GAL1*-luciferase induction profiles of the indicated yeast strains after a previous galactose encounter (+ memory) or in naïve cells (-memory). In all experiments, at least 3 independent biological replicates were analyzed.

One key factor for a more sensitive galactose recognition and signaling is the expression of the galactose sensor Gal3 (19, 28). Therefore, we analyzed the dose response behavior of the *GAL3* upstream control regions of *S. bayanus* in comparison to the *S. cerevisiae* BY4741 reference. We additionally included the *GAL3* promoter variants from suboptimal galactose consumers, such as *S. cerevisiae* Y12 and DBVPG6044, in this study. We recorded the complete dose-response profiles of all *GAL3* promoter variants by time elapsed luciferase assays in the BY4741 genetic background (Figure 5A). We first noticed that *GAL3p*_*S*.*bayanus*_ displayed a significantly higher basal activity as compared to all other *GAL3* promoters (Figure 5B). Moreover, *GAL3p*_*S*.*bayanus*_ was activated by very low galactose concentrations (<0.1%) to almost complete efficiency (Figure 5C). Thus, the Gal3 expression in *S. bayanus* is driven by a constitutively more active promoter, which is more sensitively activated by low inducer concentrations. Inspection of the respective nucleotide sequences of the *GAL3* upstream control regions revealed no obvious changes within the perfectly conserved binding region for the Gal4 transcriptional activator (Figure 5D). However, we observed important differences in the sequences of two Mig1 repressor binding sites in the *GAL3* upstream sequences. The reference GAL3p_BY4741_ contains two predicted consensus sequences for Mig1, proximal site 1 between the Gal4_UAS_ and the transcription start site and distal site 2 just upstream of Gal4_UAS_ (Figure 5D). Site 1 perfectly matches the previously characterized Mig1 consensus with the 3’-G/C box and the adjacent 5’-AT-rich flanking region (38). Site 2 only conserves the G/C box and might therefore not be functional. In any case, the *GAL3p*_*S*.*bayanus*_ sequence, but not the other *S. cerevisiae* variants, shows several point mutations in the essential G/C box motifs, thereby inactivating both Mig1 binding sites (Figure 5D). These data suggest that *S. bayanus* has evolved an especially sensitive galactose consumption behavior, which at least in part, might be caused by its derepressed and hyper-activatable *GAL3* control lacking Mig1 repression.

**Fig. 5.**
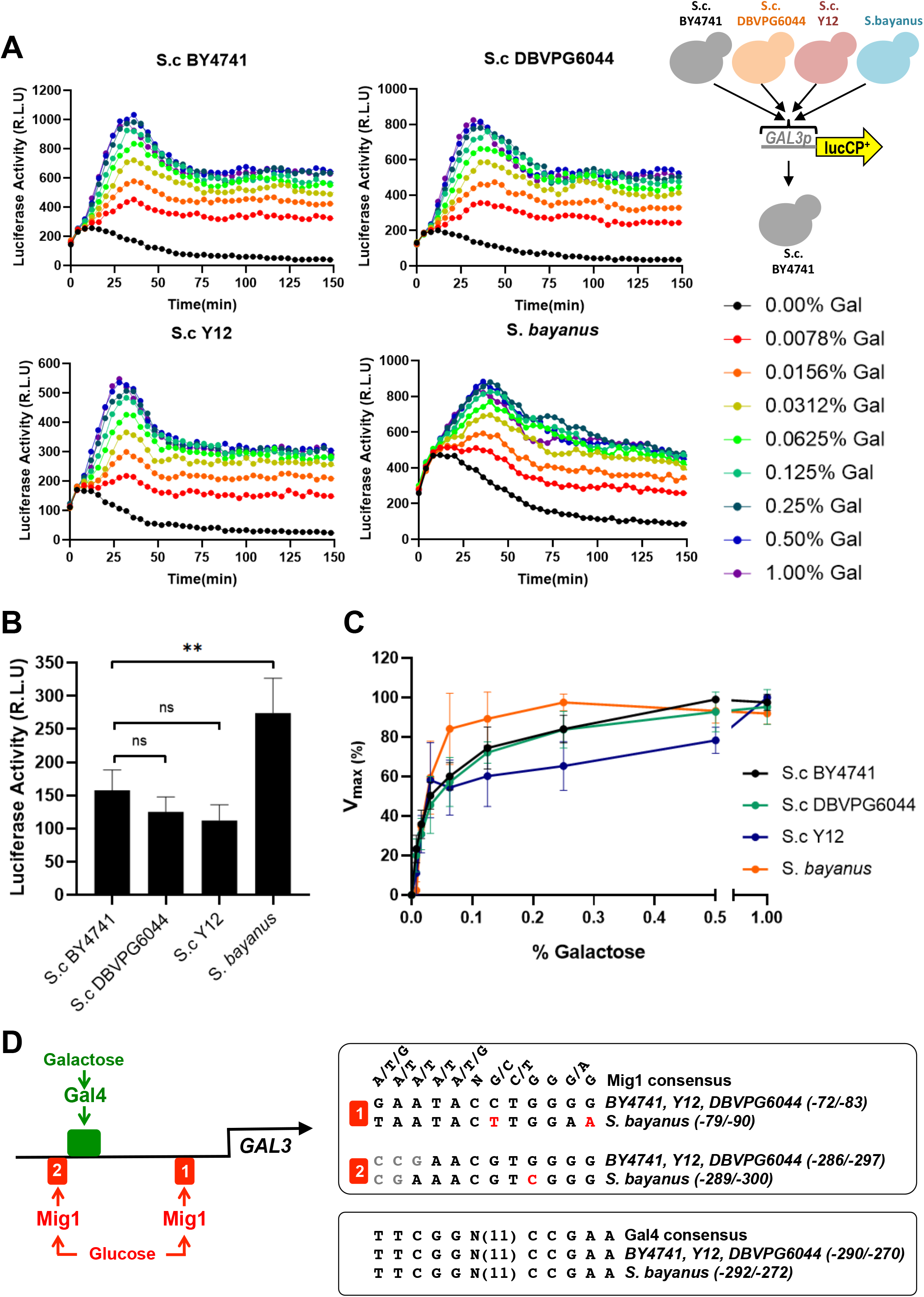
*S. bayanus* contains a constitutively active and galactose hypersensitive *GAL3* promoter. (A) Dose-response profiles of different *S. cerevisiae* and *S. bayanus GAL3* variant-luciferase reporters. Live cell luciferase reporter assays were performed in *S. cerevisiae* BY4741 containing the indicated *GAL3p* variants. Results are depicted as medium values from at least 3 biological replicates. (B) Representation of the basal *GAL3* promoter activity of the indicated yeast strains. Data are mean (n=3) +/- SD. Statistically significant differences were determined with the unpaired Student’s t-test (** p<0.01). (C) Comparison of the sensitivity of *GAL3* induction across the different variants. Representation of the maximal light increment (v_max_) versus the galactose inducer concentration for each yeast strain. v_max_ values were set to 100% for the inducer concentration, which leads to maximal induction. (D) *GAL3* promoter sequence comparison. Highlighted are the unique Gal4 UAS motif and the proximal (1) and distal (2) Mig1 repressor binding sites. Nucleotide substitutions in *S. bayanus*, which inactivate the Mig1 sites, are indicated in red.

### Specialization towards desensitized galactose consumption: the case of *S. cerevisiae* Y12

Our previous results showed that the *S. cerevisiae* Y12 isolate needed higher galactose concentrations in the medium to efficiently consume the alternative sugar. This behavior is characterized by a less sensitive *GAL1* gene induction, a pronounced diauxic lag phase and decreased galactose growth rates. We investigated in depth this sugar consumption strategy as an example of a naturally evolved enhanced sugar discrimination. We first looked in detail at the *GAL1* induction capacity of Y12 and found that it was impaired only at very low galactose concentrations showing a significant response delay (Figure 6A). High galactose concentrations, however, caused a timely and even more dynamic *GAL1* activation in Y12 as compared to BY4741. This more discriminate galactose behavior was perfectly recapitulated by the absolute induction velocities, which were lower than BY4741 in a range up to 0.02% galactose and actually higher than BY4741 for galactose >0.02% (Figure 6B). We next tested whether this apparent less affinity for low galactose doses had consequences for the susceptibility to the toxic glucose analogue GlcN. Indeed, we found a pronounced sensitivity of *S. cerevisiae* Y12 to low GlcN doses, indicating a higher preference of Y12 for glucose consumption in the presence of alternative galactose (Figure 6C). At low inducer concentrations, the yeast *GAL* response can be bimodal with only a fraction of the cells in the population actively engaged in gene expression. In order to determine the fraction of actively responding cells during the transition from the off to the on state, we measured the *GAL1p*-driven GFP expression upon low and high galactose induction by flow cytometry (Figure 6D). We found that high galactose concentrations caused a rapid and comparable transition to GFP positive cells in BY4741 and Y12. As expected, the transition from the off to the on state occurred more slowly in BY4741 upon low galactose induction (<0.02%). However, the Y12 isolate showed a severe induction delay upon those inducer concentrations (Figure 6D). The comparison of the induction levels in both yeast strains again demonstrated that Y12 cells discriminate much more low and high galactose concentrations in the medium (Figure 6D). We finally confirmed that, like the BY4741 reference strain, Y12 cells possess a strong positive memory upon repeated galactose encounters (Figure 6E). However, even in galactose experienced Y12 cells, it seemed that low galactose stimulation still caused an inefficient response as compared to the very much improved induction in experienced BY4741 cells (Figure 6E). This observation suggested that Y12 cells have a less sensitive Gal3 sensor, which needs higher galactose levels to efficiently induce *GAL* gene expression. In order to test this hypothesis, we expressed the Gal3_BY4741_ and Gal3_Y12_ sensor variants from the strong *GAL1*_BY4741_ promoter and compared their dose dependent activation of a *GAL1p*-luciferase reporter (Figure 7A, B). Transactivation of both Gal3 sensors was indistinguishable at high galactose concentrations, while low galactose doses triggered a more efficient *GAL1* induction through Gal3_BY4741_ as compared to Gal3_Y12_ (Figure 7C). These data demonstrate that *S. cerevisiae* Y12 possesses a less sensitive Gal3 sensor. Indeed, the concentration dependent Gal3_Y12_ activation profile in BY4741 cells was similar to the one observed in Y12 galactose experienced cells (Figure 7D), indicating that the desensitized galactose signaling of the Y12 strain stems mainly from its less sensitive Gal3 sensor protein.

**Fig. 6.**
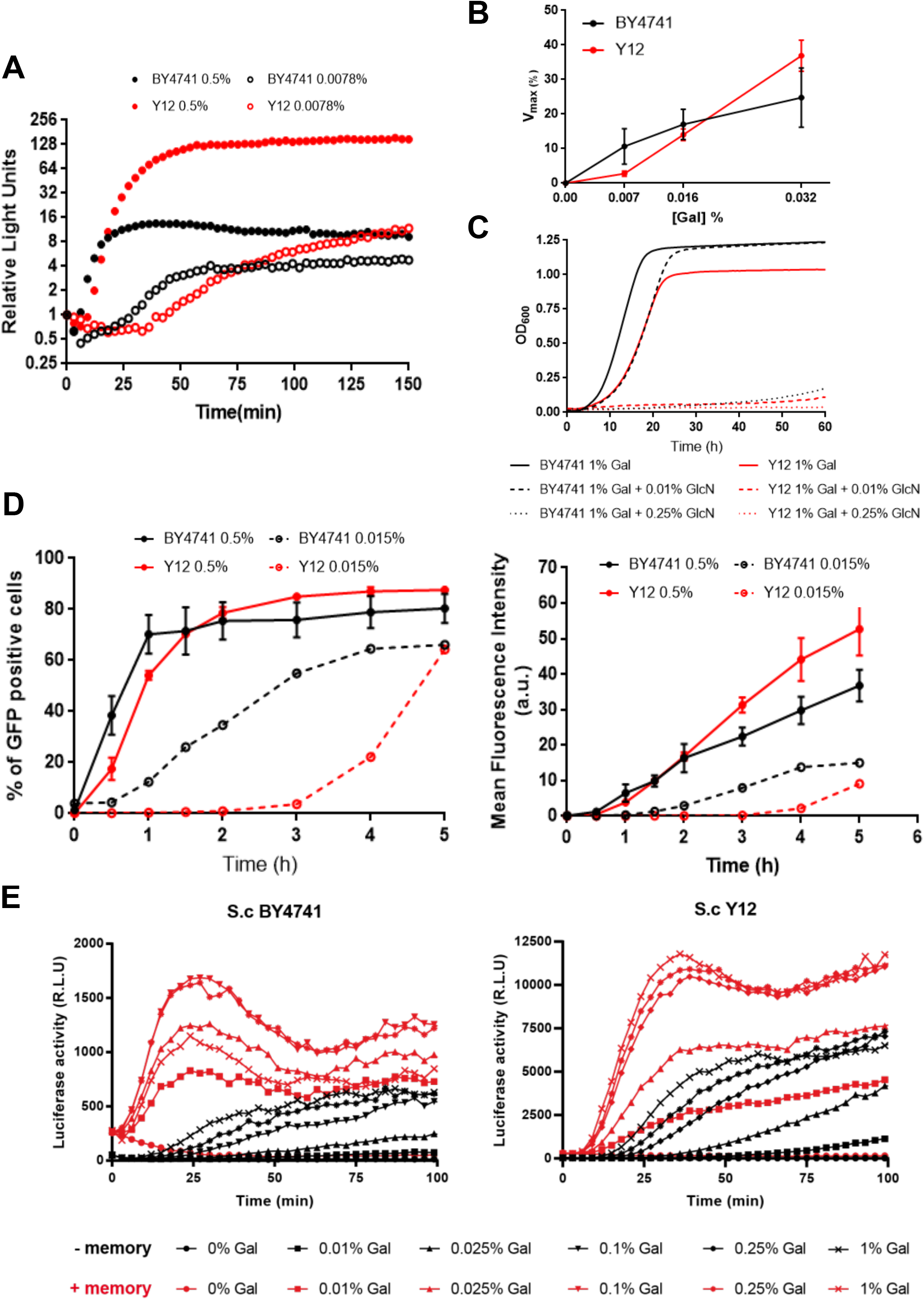
Features of a desensitized galactose consumption strategy in *Saccharomyces cerevisiae* Y12. (A) The relative luciferase expression of a *GAL1*-luciferase reporter is compared between *S. cerevisiae* BY4741 and Y12 for the indicated inducer concentrations. Data are taken from the experiment described in Figure 1B, C. (B) The relative light increment (v_max_) of a *GAL1*-luciferase reporter is compared between *S. cerevisiae* BY4741 and Y12 for the indicated inducer concentrations. Data are taken from the experiment described in Figure 1B, C. (C) Resistance to the glucose analogue glucosamine. Yeast cells were grown in pure galactose medium (1%) with or without the addition of the indicated amounts of glucosamine (GlcN). (D) Flow cytometry analysis of the *GAL1-GFP* induction process in *S. cerevisiae* BY4741 and Y12 upon high (0.5%) and low (0.015%) inducer concentrations. Left panel: Comparison of the fraction of GFP positive cells; right panel: comparison of the induction levels of the GFP positive cells. (E) *S. cerevisiae* Y12 shows a transcriptional galactose memory. Comparison of the *GAL1*-luciferase induction profiles of the indicated yeast strains after a previous galactose encounter (+ memory) or in naïve cells (-memory) as described in Figure 4D. In all experiments, at least 3 independent biological replicates were analyzed.

**Fig. 7.**
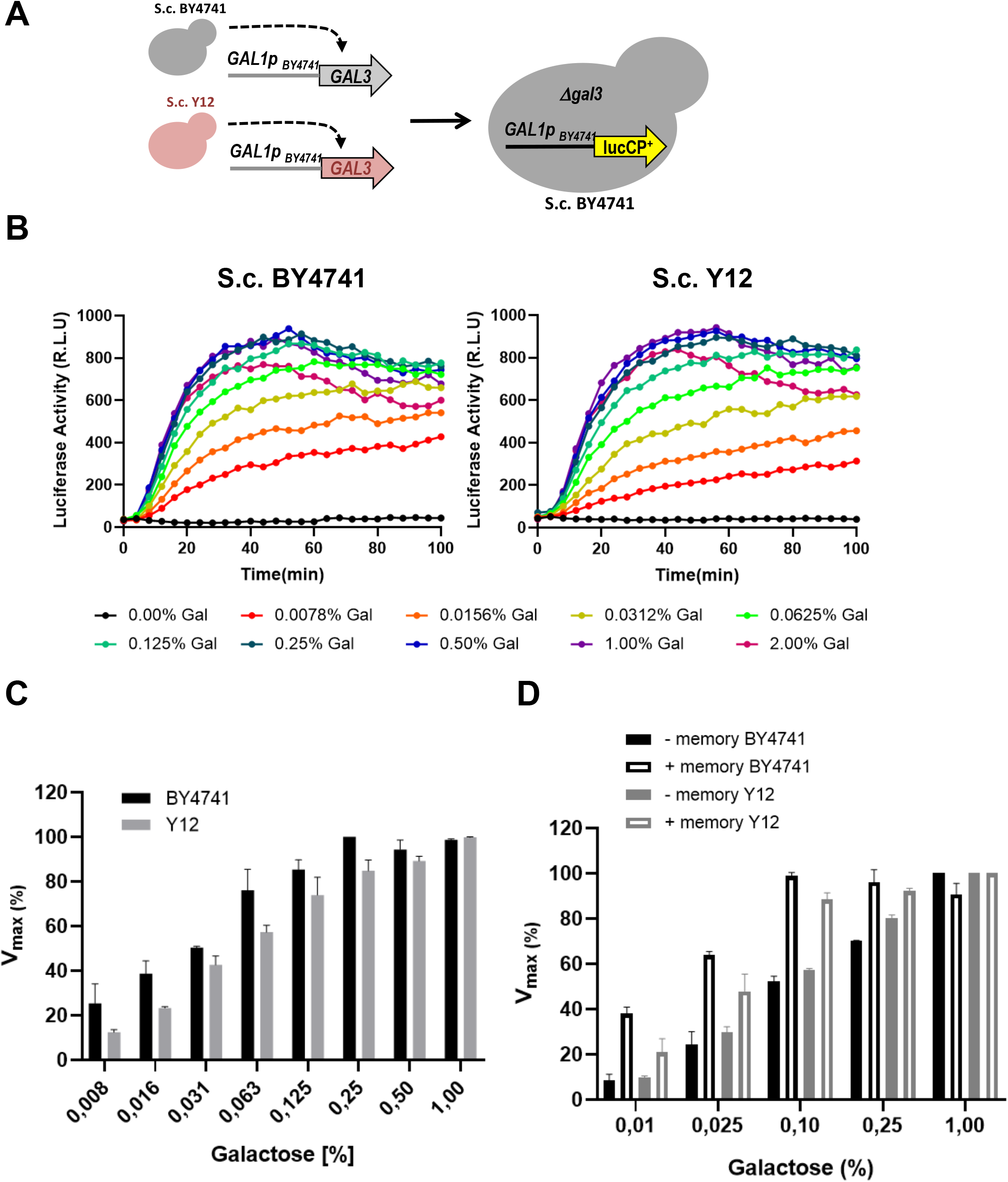
*S. cerevisiae* Y12 harbors a less sensitive Gal3 sensor. (A) Experimental strategy to quantitatively compare the trans-activation capacities of the galactose sensors Gal3_BY4741_ and Gal3_Y12_. (B) Dose response profiles of *GAL1p*-luciferase through Gal3_BY4741_ and Gal3_Y12_. (C) Comparison of the sensitivity of *GAL1* induction. Representation of the maximal light increment (v_max_) versus the galactose inducer concentration for each Gal3 variant. V_max_ values were set to 100% for the inducer concentration, which leads to maximal induction. (D) Gal3_Y12_ confers less sensitive *GAL* gene induction even after previous galactose encounter. Data are taken from the experiment described in Figure 6E and represented as in (C). In all experiments, at least 3 independent biological replicates were analyzed.

### Extremely insensitive galactose consumption: the case of the West African *S. cerevisiae* DBVPG6044 and NCYC110

The West African yeast isolates investigated here displayed an extraordinarily long lag phase and poor, but not absent, growth rates on galactose media. Accordingly, no *GAL1* gene induction was detected in the first hours of galactose stimulation. Additionally, short term memory effects, such as previously found here for *S. cerevisiae* BY4741 or Y12, were completely absent (data not shown). The DBVPG6044 and NCYC110 isolates required a previous galactose pre-incubation for several days in order to improve galactose growth parameters (Figure 3) or *GAL1* gene induction (data not shown). We re-sequenced the *GAL3*_DBVPG6044_ allele and found a single nucleotide substitution, C_456_ to A_456_, which created a premature stop codon after aa_151_ (Figure 8A). This mutation has been previously reported for some West African yeast isolates including DBVPG6044 and NCYC110 (39). However, if this inactivation of Gal3 function was the only defect in the *GAL* system in these isolates, their galactose consumption should be restored by re-introducing a functional Gal3 wild type copy. Indeed, when we transformed DBVPG6044 or NCYC110 yeast cells with a *GAL3*_BY4741_ allele, both strains recovered robust growth on pure galactose media (Figure 8B). These data indicated that the West African isolates DBVPG6044 and NCYC110 have adopted a slow and insensitive galactose response by genetic interruption of their *GAL3* sugar sensor.

**Fig. 8.**
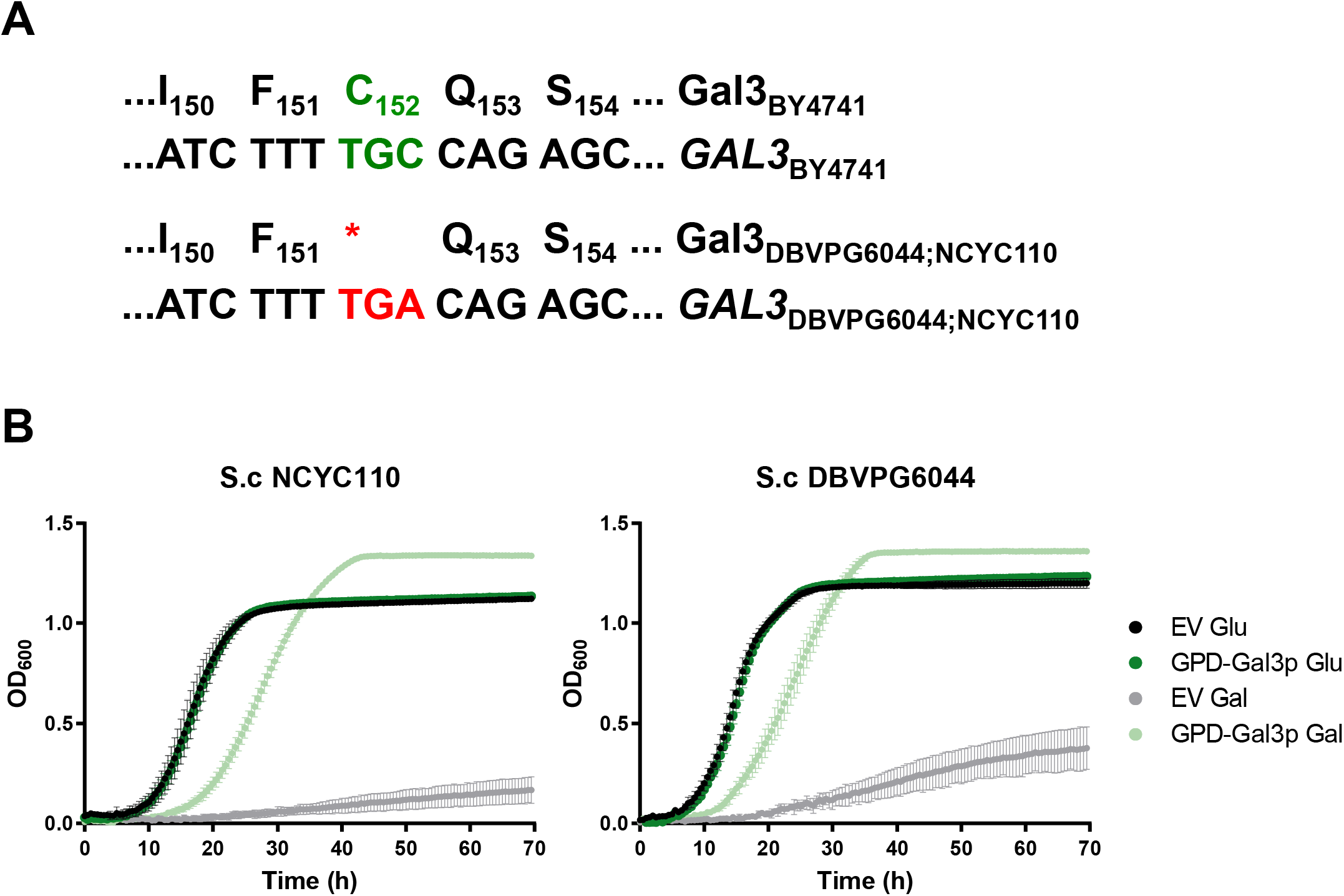
West African *S. cerevisiae* isolates DBVPG6044 and NCYC110 are poor galactose consumers due to an inactivated *GAL3* allele. (A) Sequence comparison of *GAL3* of the indicated *S. cerevisiae* isolates. (B) Re-introduction of a Gal3_BY4741_ copy restores efficient galactose growth of *S. cerevisiae* DBVPG6044 and NCYC110. Cells harboring the empty plasmid control (EV) or the GPD-GAL3_BY4741_ expression plasmid were grown in synthetic glucose or galactose media. Data are mean (n=3) +/- SD.

## Discussion

The natural habitats of microorganisms often experience fluctuations in the nutrient composition. These changes trigger the induction of the metabolic pathways needed to efficiently utilize the available nutrients (2, 40). Related microbial organisms have evolved divergent strategies, both in terms of the nutrients recognized and of the efficiency of metabolizing them (39, 41). In general, the decision to utilize a particular alternative nutrient is conditioned by the cost to maintain the metabolic enzymes even when the nutrient is scarce and the benefit for cellular energy metabolism by efficiently consuming the metabolite (20, 42, 43). Here, we reasoned that in the case of galactose utilization in yeast, different sugar preferences should be quantifiable by comparing the dose-response induction profile of its principal metabolic enzyme Gal1. The destabilized luciferase technology applied by us faithfully identifies such changes in the *GAL* dose-response across different *S. cerevisiae* natural isolates and related *Saccharomyces* species.

In all yeasts analyzed here, *GAL1* gene expression is repressed in the absence of galactose. Thus, divergent galactose preferences should arise from a different facility to overcome this repression and efficiently transcribe *GAL1*. The key *GAL* inducer is the *GAL3* sensor, which arose from the original *GAL1* galactokinase gene by gene duplication. Further specialization of *GAL3* through evolution turned the protein from a galactose phosphorylating enzyme to a highly sensitive galactose sensor needed to switch the Gal4-Gal80 complex to an efficient transcriptional activator of *GAL1*(44–46). Importantly, *GAL3* expression itself is galactose dependent (47), thus the sensitivity of the *Saccharomyces GAL* switch could be either modulated by the expression dynamics or the intrinsic activity of a single sugar sensor, Gal3. Indeed, the dominant role of polymorphisms in the *GAL3* locus in the diversification of galactose responses in different *S. cerevisiae* isolates has been recently demonstrated (19). Here we identify very divergent galactose consumption strategies, which at least in great part, can be explained by differential expression or altered sensitivities of the Gal3 sensor.

The galactose behavior of *S. bayanus* is an extreme adaptation towards highly efficient galactose consumption, while keeping the *GAL* system still under control of galactose induction. This induction, however, is very much improved, so that maximal *GAL* gene induction is achieved in this yeast at 5 to 10-fold less galactose concentrations as compared to its *S. cerevisiae* relatives. Consequently, this hypersensitive *GAL* switch enables a continuous proliferation in habitats where glucose is exhausted in the presence of galactose. Here, *S. bayanus* outgrows other yeast species with a diauxic lag phase. Thus, the *S. bayanus* galactose consumption strategy might be advantageous in rapidly changing mixed sugar environments. One key event in creating this extra sensitive *GAL* switch is the here reported modification of the *GAL3* upstream control region. An increased basal level and more sensitive galactose induction of *GAL3* in *S. bayanus* seems to create the hyper sensitive *GAL* response. It is likely that the Gal3 levels in *S. bayanus* are not rate limiting even in the absence of the alternative sugar. This would explain the complete lack of transcriptional memory in this yeast. Improvement of the *GAL* switch after repeated galactose exposures depends on higher Gal3 (and to a lesser extent Gal1) protein levels (31), and even slight overexpression of Gal3 strongly improves the sensitivity of *GAL* induction in laboratory yeast strains (28, 48). Consistent with our finding, it has been reported previously that *GAL* upstream control regions are the key elements that confer delayed galactose consumption in *S. cerevisiae* as compared to the long-diverged *S. bayanus* (49). Specifically, the single substitution of the *GAL3* promoter in *S. bayanus* with the *S. cerevisiae* counterpart restored an evident diauxic lag (49), which confirms that Gal3 expression levels are key to *GAL* switch sensitivity and galactose selectivity.

Diversification of the *GAL* switch among different *S. cerevisiae* isolates dominantly depends on evolutionary differences in the *GAL3* locus (19). However, how specific mutations in the Gal3 protein confer different *GAL* switch characteristics remained unknown. The *S. cerevisiae* Y12 behavior is characterized here as an intra-species adaptation towards higher galactose discrimination. Here, the basis is not an altered Gal3 expression, as the *GAL3*_Y12_ promoter is equally active in the absence or presence of low galactose concentrations. The Gal3_Y12_ sensor needs higher galactose concentrations to be fully active and operate the *GAL* switch through Gal80 inactivation. Several polymorphisms are present in the Gal3_Y12_ coding sequence: K_122_R, I_135_V, P_137_L, Q_149_L, L_302_P, H_352_D, and L_370_P. The reversible Gal3 interaction with the Gal80 repressor requires binding of ATP and galactose (16, 18). Thus, some amino acid substitutions in Gal3_Y12_ may have specific effects on its galactose/ATP or Gal80 binding efficiency. The Gal3_Y12_ sensor is an interesting candidate for further characterization by biochemical experiments. Less sensitive galactose recognition might be advantageous in environments, where galactose is rarely available or at concentrations, which do not sustain cell proliferation. Y12 cells have tuned down the Gal3 sensitivity and thus raised the galactose threshold needed to activate the *GAL* switch and to metabolize the alternative sugar only at higher concentrations. A further adaptation towards even less sensitive galactose consumption has occurred in some West African *S. cerevisiae* isolates by a premature stop in the *GAL3* coding region. Those yeast isolates have been previously categorized as galactose non-fermenters (19, 39). However, we show here that these yeasts have not given up metabolizing galactose, instead they have evolved an extremely insensitive *GAL* switch by inactivating their Gal3 sensor. This likely adaptation to very infrequent galactose availability still enables to consume the alternative sugar at very low rates and to display a positive memory after a prolonged galactose encounter presumably via the remaining Gal1 galactokinase (24, 30). In summary, the function of a dedicated and inducible nutrient sensor in the *GAL* switch has enabled the evolution of very divergent galactose consumption strategies in yeast.

## Acknowledgments

The authors thank Jun-Yi Leu for the kind gift of *S. cerevisiae* natural isolates and *Saccharomyces bayanus, Saccharomyces mikatae* and *Saccharomyces paradoxus* strains.

